# Sequential Effects in Reaching Reveal Efficient Coding in Motor Planning

**DOI:** 10.1101/2024.09.30.615975

**Authors:** Tianhe Wang, Yifan Fang, David Whitney

## Abstract

The nervous system utilizes prior information to enhance the accuracy of perception and action. Prevailing models of motor control emphasize Bayesian models, which suggest that the system adjusts the current motor plan by integrating information from previous observations. While Bayesian integration has been extensively examined, those studies usually applied a highly stable and predictable environment. In contrast, in many real-life situations, motor goals change rapidly over time in a relatively unpredictable way, leaving it unclear whether Bayesian integration is useful in those natural environments. An alternative model that leverages prior information to improve performance is efficient coding, which suggests that the motor system maximizes the accuracy by dynamically tuning the allocation of the encoding resources based on environmental statistics. To investigate whether this adaptive mechanism operates in motor planning, we employed center-out reaching tasks with motor goals changing in a relatively unpredictable way, where Bayesian and efficient coding models predict opposite sequential effects. Consistent with the efficient coding model, we found that current movements were biased in the opposite direction of previous movements. These repulsive biases were amplified by intrinsic motor variability. Moreover, movement variability decreased when successive reaches were similar to each other. Together, these effects support the presence of efficient coding in motor planning, a novel mechanism with which the motor system maintains flexibility and high accuracy in dynamic environments.

## Introduction

Our experiences shape how we perceive and interact with the world. For instance, when facing ambiguous visual stimuli, humans are more likely to identify familiar objects than novel ones^1,2^. Similarly, in reaching tasks, participants infer the state of the environment based on previous observations to apply an optimal action^3–5^. Those processes have been formalized with Bayesian observer models^2,3,6,7^, which posit that the system maintains an internal prior based on past observations and integrates this prior knowledge with new sensory input to form optimal estimates^8^. This integration reduces variability in estimation, thereby increasing overall response accuracy.

The Bayesian framework has significantly influenced theories of motor control and motor learning^3,5,9,10^. For example, when localizing hand position, agents combine information from different sensory inputs in a Bayesian-optimal way^9,11,12^. Moreover, sensorimotor learning theories posit that the system uses error information to update the sensorimotor map based on a Bayes rule^13,14^. Furthermore, Bayesian principles have been applied to interpret history effects in reaching tasks. Previous research has shown that after repeatedly performing one action, future movements would be biased toward the repeated direction—a phenomenon known as use-dependent learning^15,16^. The Bayesian model explains this phenomenon by positing that the integration of previous motor goals with the current goal induces a behavioral bias toward the prior mean^17^ (Fig 1a-b).

**Figure 1.**
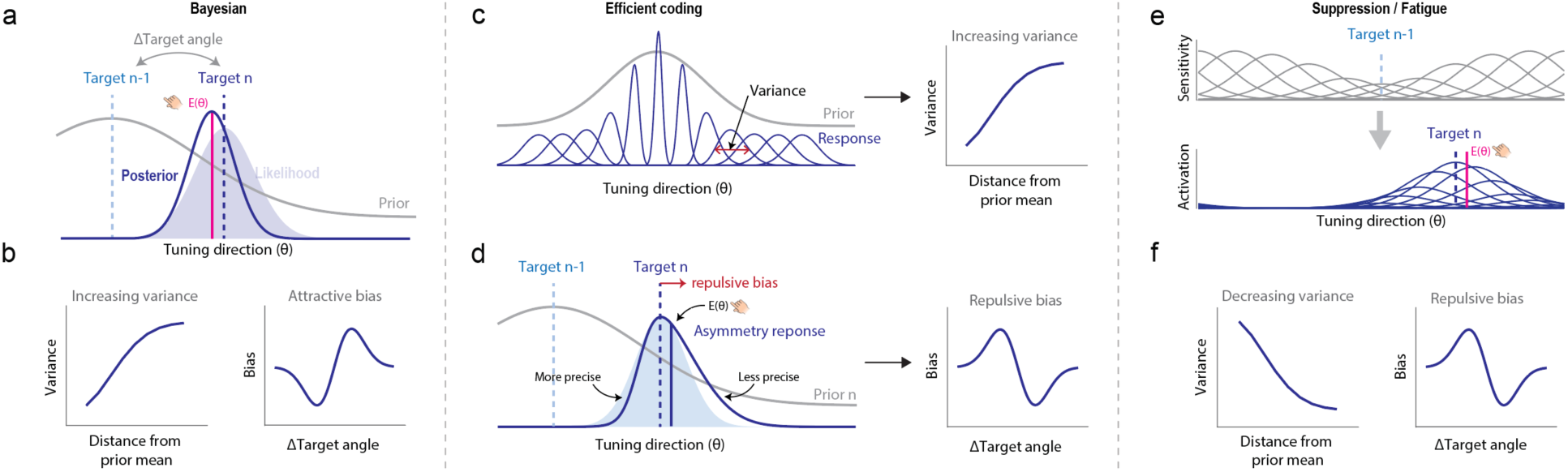
Models for sequential effects in motor planning. a) Illustration of a Bayesian model in motor planning. b) The Bayesian model predicts increasing variance and attractive bias. c) Illustration of an efficient coding framework. The motor system encodes a motor command based on the perceived target following an efficient coding rule. Specifically, the precision of encoding the direction of movement should match the prior. The blue curves indicate the encoding function to different movement directions, with its variability constrained by the prior distribution. The variability in movement (inaccuracy) increases as the current movement diverges from the prior mean. d) Given the heterogeneity of encoding resources across the space, the encoding function for a given movement direction is asymmetric, with a flatter tail extending away from the prior (see method). Intuitively, this is because the encoding variability is smaller on the prior side and larger on the opposite side. E(θ) represents the mean of this distribution, which is biased in the direction opposite to the prior mean. The shaded blue distribution illustrates the response in the absence of efficient coding. In this context, we assume the prior is primarily influenced by the movement direction in the previous (n-1) trial. As such, the efficient coding model predicts a repulsive sequential bias. e) Illustration of the suppression model. The sensitivity of the unit tuned to the previous movement decreases, which causes an asymmetric representation of the subsequent movement direction. f) The suppression model predicts decreasing variance and repulsive bias.

While Bayesian models have been extensively tested in motor control, these tasks often share two key characteristics that may limit the generalizability of those findings. First, the targets used in previous studies frequently possess strong statistical structures, making it easy for participants to anticipate future movements^18^. For example, many studies have sampled targets from unimodal distributions^15,17,19^, allowing participants to predict the mean movement direction. Even in center-out reaching tasks, most experiments have utilized only a very small number of targets^20–23^, thereby confining the possible motor plans to a very limited subspace. Second, those studies examined effects based on a stable representation of the prior, which is estimated from many observations and remains constant over time.

However, contrary to those two prerequisites, in many real-life situations, motor goals can change rapidly across time in a relatively unpredictable way^18,24^. For example, a basketball player needs to constantly change their motor plans in real-time—suddenly shifting direction, accelerating, jumping, or making quick passes—in response to the unpredictable actions of opponents and teammates. Similarly, when shopping for groceries, the location of each item on the shelf varies, so reaching for one product is independent of where the next one might be. In these cases, the motor system must plan each movement independently rather than relying on a consistent prior. It remains unclear whether Bayesian mechanisms would be beneficial in such volatile environments. Alternatively, other computational mechanisms might be applied when motor goals vary on a short timescale.

An alternative framework, known as efficient coding, offers a distinct perspective on how the brain is constrained by environmental statistics^25,26^. Efficient coding posits that the nervous system minimizes redundancy and maximizes the use of available resources (Fig 1c)^27–30^. In the perceptual domain, it refers to a strategy to encode information in the most compact and effective manner^25,26^. In motor control, efficient coding can be taken as a way to avoid redundancy in the control system and maximize overall accuracy. For example, rather than controlling each muscle individually, the brain coordinates groups of muscles into synergies or modules^31,32^. Moreover, the motor system may dynamically allocate encoding resources based on environmental statistics to maximize the mutual information it carries (Fig 1c). Interestingly, previous theoretical work^27,28^ suggested that this process will induce an encoding bias in the opposite direction of the prior mean (Fig 1d), a prediction opposite to the Bayesian model. While this principle has been extensively studied in perceptual systems^25,27,33,34^, it is less clear whether it applies to motor planning.

To understand how motor planning is adjusted based on prior information in volatile environments, we aim to examine sequential effects in a center-out reaching task where the target is randomly sampled from a uniform circular distribution. As such, no mean movement direction can be predicted, and each movement is completely independent. While the overall prior is not informative in this situation, the system may maintain a short-term prior that is updated quickly when the goals change rapidly^35,36^. This flexibility could be particularly beneficial for the motor system, which may need to repeat similar movements in quick succession, though not necessarily over extended periods.

Following the Bayesian model, prior updating would generate an attractive sequential effect, where the current response is biased toward previous responses that have just been performed (Fig 1a-b)^17,35,36^. This effect, known as serial dependence, has been widely observed in perceptual tasks across different modalities^37–40^. On the other hand, the efficient coding models predicts the current movement will be biased away from the previous movement. As such, efficient coding and Bayesian integration generate opposite predictions on the sequential effect, allowing us to distinguish between these two theoretical frameworks in the current study. Additionally, we explored the temporal dynamics of these sequential effects and how they are modulated by intrinsic motor variability, which helps further differentiate between the models. The empirical and computational results together illustrate how the motor planning system adapts to the statistical properties in a dynamic environment.

## Results

### A repulsive sequential bias in reaching

To examine motor sequential effects in an unpredictable environment, in Exp 1, we employed an online reaching experiment where participants control a cursor with a trackpad. In each trial, a target appeared at a random position along an invisible circle and participants made a center-out reaching movement from a central start position towards the target (Fig 2a). The cursor was invisible during the movement to prevent online corrections. Endpoint feedback was provided after the movement to indicate the hand position at the target radius.

**Figure 2.**
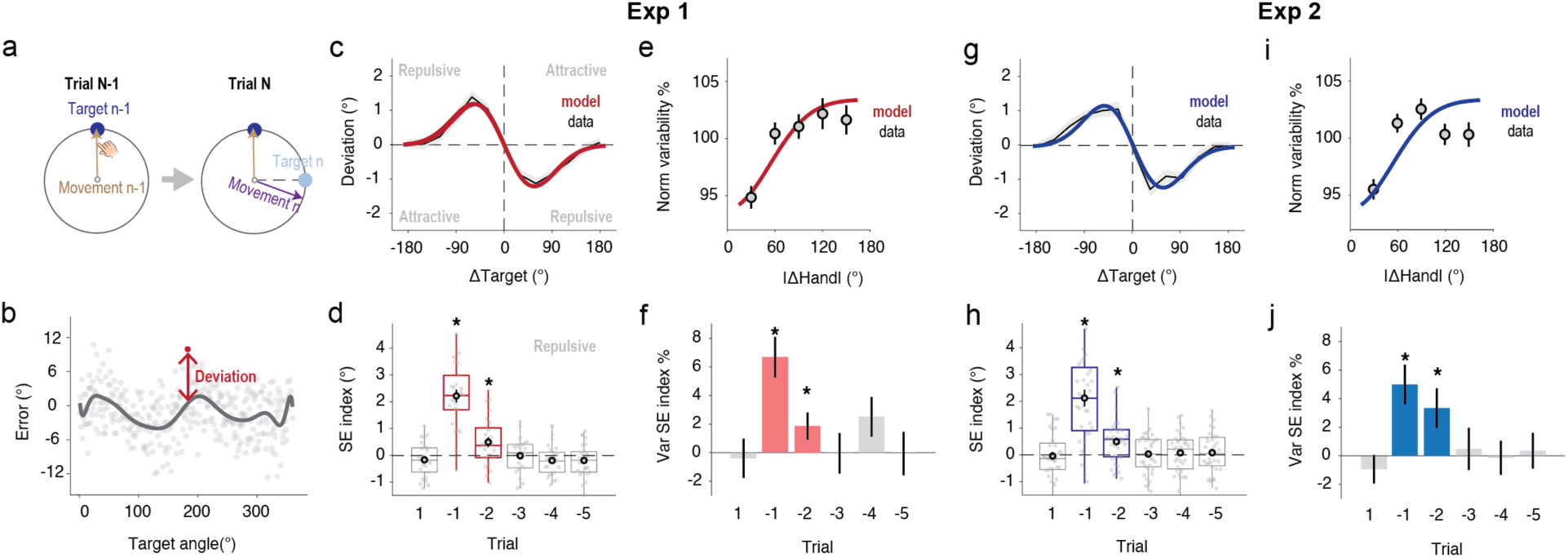
Sequential effects in reaching. a) An illustration of the repulsive bias predicted by the efficient coding model. b) The motor bias of a sample participant. The gray curve shows the smoothed motor bias function. The deviation was defined as the difference between the current response and the bias. d) Sequential bias in Exp 1: The motor deviation of trial n was plotted as a function of the difference in the target angle between trial N and trial N- 1 (Δtarget). This figure shows a repulsive sequential effect. e) SE index showing the influence of the target in trial N+1, and trials N-1 to N-5 on the direction of reaching in trial n. f) The normalized movement variance increased with the distance between movement N and the movement N-1 (|Δhand|). g) Sequential effect (SE) index derived from a general linear model between normalized movement variance and |Δhand| for trial N+1, and trials N-1 to N-5. h-k). Sequential effects in Exp 2. The only difference between Exp 1 and 2 was the lack of feedback in Exp 2. Results were very similar to panel c-f. In c, e, g, i, the black curve indicates data, and the colored curve indicates the prediction of the best-fitted efficient coding model (see method). Error bars and shaded areas indicate standard error. *, p<0.02.

To obtain an accurate assessment of the sequential effect, we eliminated the impact of systematic bias by fitting a motor bias function based on the target position (Fig S2) for each participant. The residual error, representing the deviation from this fitted motor bias function (Fig 2b), was defined as the deviation^35,36,41,42^ and was used to analyze the sequential effects.

We observed a repulsive sequential bias in reaching. When movement deviation is plotted as a function of the angular difference between the current and previous targets (ΔTarget). We found that the direction of movement in the current trial (trial n) was biased away from the previous target (trial n-1, Fig 2c), with the magnitude of this repulsive effect increasing with ΔTarget, peaking at 1.5° for a ΔTarget of approximately 60°. This bias function aligns with predictions from the efficient coding model rather than the Bayesian model. To quantify the size of the sequential bias, we calculated a sequential effect (SE) index, defined as the difference between the average bias for Δ Target from −90° to 0° and 0° to 90°. A positive SE index signified a repulsive effect. We found an SE index significantly larger than 0 in trial N-1 (*t*(25)=9.5, *p*<.001, d=1.9) and trial N-2 (*t*(25)=2.8, *p*=0.010, d=0.55), but not in trial N-3 (*t*(25)=-0.10, *p*=0.92, d=-0.02; Fig 2d). We further showed that the attenuation of the sequential effects depended on both passing time and intervening information by manipulating the inter-trial-intervals in Exp S1 (See Supplementary Result 1, Fig S1).

### Sequential effects in reaching align with the efficient coding model

The repulsive sequential biases observed in Exp 1 support the predictions of the efficient coding model rather than the Bayesian model. However, we note that an alternative explanation for this effect could be suppressive modulation or fatigue affecting the units tuned to the recently performed movement direction. The suppression model predicts a repulsive sequential bias due to the asymmetric responses of units on the suppressed and unsuppressed sides (Fig. 1e). However, unlike the efficient coding and Bayesian models, which both facilitate the processing of repeated stimuli, the suppression model posits that movements closer to the previous movement are encoded with less precision and exhibit greater variability compared to those further away (Fig. 1f).

To compare the efficient coding model and the suppression model, we measured the sequential effect as a function of movement variance. Consistent with the efficient coding model, the movement variance increased with the distance between the directions of two consecutive movements (Fig 2g–h). As such, the repulsive sequential bias is unlikely to be due to fatigue. We quantified this effect by fitting a sigmoid function with motor variance as the dependent variable and the difference in hand angle as the independent variable, pooling data from all participants. The SE index was defined as the amplitude of the sigmoid predicted by the best-fit function (Fig 2e). Similar to the sequential bias, this SE index of movement variance dropped rapidly across trials: a significant effect was found for only trial N-1 (p < 0.001) and N-2 (p = 0.026, Fig 2f), as measured by bootstrap resampling. The effect disappeared for trial N-3 (p = 0.98), which occurred ∼6s prior. The sequential effect in movement variance confirms that, while there was a repulsive bias, the motor system utilizes previous information to increase overall accuracy in reaching, consistent with the key idea of the efficient coding model.

We note that a potential issue in Exp 1 is that endpoint feedback could facilitate the recalibration of the sensorimotor system. This recalibration process might be specific to areas near the recently reached positions, potentially contributing to the observed sequential effect in motor variance. To eliminate the potential influence of sensorimotor recalibration, we conducted a replication of Exp 1 without endpoint feedback (Exp 2). The results showed remarkably similar sequential effects in both movement bias and variance (Fig 2g-j), confirming that these effects were independent of visual feedback or sensorimotor recalibration.

### Dissociating sequential effect in movement and perception

Sequential effects have been widely observed in perceptual tasks^37,38,43^. While we tried to minimize the role of visual uncertainty and perceptual working memory in Exp 1, a more rigorous examination was necessary to differentiate effects originating in the motor versus the perceptual systems^44^. To this end, in Exp 3 we implemented a design where participants were instructed to either move directly towards the target (Standard condition, 75% of trials) or in the opposite direction (Opposite condition, 25% of trials). As a critical test of the source of these sequential effects in Exp 1, we analyzed scenarios where trial N-1 was an Opposite trial and trial N was a Standard trial (Fig 3a, bottom). In such cases, the inconsistency between movement direction and target in trial N allowed us to disentangle the influences of target location representation from the motor control itself. If the repulsive bias was primarily driven by motor factors, the direction in trial N would be repelled by the movement in trial N-1. Conversely, if perception of the target location was the cause, trial N’s reaching direction would be repelled by the perceived location of the target in trial N-1.

**Figure 3.**
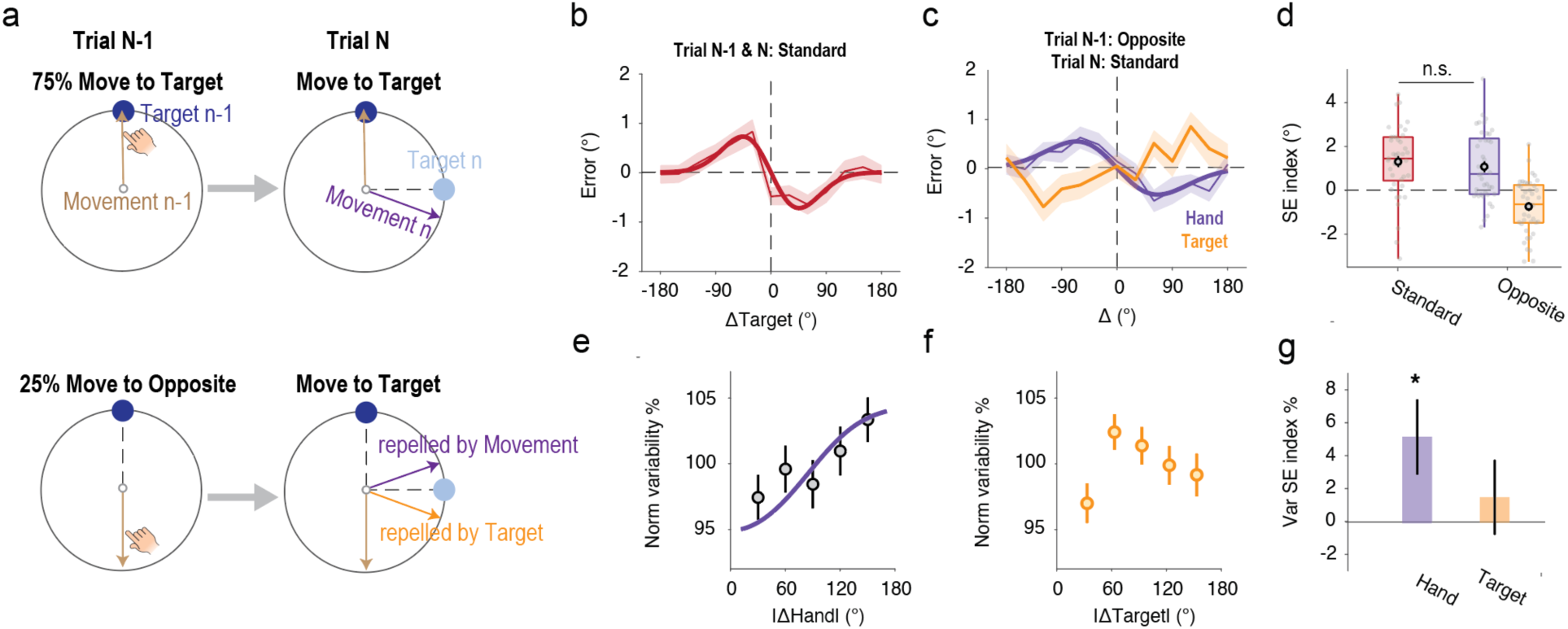
Sequential effects in reaching were associated with motor movement rather than perception. a) Design of Exp 3: In 75% of trials, participants were directed to move towards the target (Standard), and in 25% of trials, they were directed to move in the opposite direction of the target (Opposite). A repulsive effect was expected when both trials N and N-1 were Standard trials (top row). In cases where trial N was a Standard trial and trial N-1 was an Opposite trial, the design allowed examination of whether the repulsive effect was triggered by target perception or movement (bottom row). b-c) The sequential effect of motor bias when both trial N and trial N-1 were Standard trials (b) or when trial N-1 was an Opposite trial and trial N was a Standard trial (c). The thin lines with shaded error bars indicate data, and the thick curve indicates the prediction of the efficient coding model. d) The sequential bias was similar across panels b and e when the SE index after the Opposite trial was measured based on Δ Hand rather than Δ Target. e-f) After an Opposite trial, movement variance increased as a function of Δ Hand (e) rather than Δ Target (f). The motor variance increased with Δ Hand rather than Δ Target. g) The coefficients of linear regression measured from panel e-f. Error bars and shaded areas indicate standard error. *, p<.001.

When two consecutive trials were both Standard trials (Fig 3a, top), we observed a repulsive bias similar to that in Exp 1 (Fig 3b). Crucially, when trial N-1 was an Opposite trial and trial N was a Standard trial, the direction in trial N was repelled away by the previous *reaching movement* (*t*(40)=4.4, *p*<.001, d=0.68), not by the previous target location (Fig 3c). The magnitude of this repulsive effect from the previous movement was comparable to that observed when both trials N and N-1 were Standard conditions, indicated by the SE indexes (*t*(40)=0.70, *p*=.49, d=0.11; Fig 3d). Moreover, when examining the sequential effect on motor variance, we found that variance increased with the difference in hand angle (*p*<.001, Bootstrap, Fig 3e) rather than the difference in target angle (*p*=0.20, Fig 3f-g). These results suggested that the sequential effects in Exp 1-2 were rooted in the motor system rather than the perceptual system.

We have shown that the sequential bias in Exp 1-3 was caused by movement; However, it remains possible that this effect could be mediated by perception. For instance, the movement from trial N-1 might repulse the perception in trial N, thereby affecting the subsequent movement (mediated hypothesis). Alternatively, the current movement could be directly repelled away by the previous movement (direct hypothesis). To distinguish between these hypotheses, we examined cases where trial N-1 was a Standard trial and trial N an Opposite trial (Fig S3a). Under the mediated hypothesis, the current perception is repelled away from the previous movement, so that we would expect the current movement to be attracted towards the movement N-1 when participants are instructed to move in a direction opposite that of target N. In contrast, the direct hypothesis predicts the current movement to be repelled from the target/movement in N-1 (Fig S3b). Our findings aligned with the direct hypothesis (Fig S3c-d). Based on the results in Exp 3, we concluded that the current movement was directly influenced by the previous movement.

We also observed a priming effect in reaction time in Exp 1-3. Consistent with the notion of efficient coding, reaction times were shorter when the current target was close to the previous target (Fig S4a-b). However, this effect may pertain more to target detection than motor planning. The reaction time for the current trial was influenced by targets presented more than 10 trials in the past (Fig S4c-d), which was very different from the temporal dynamics of sequential effects in movement direction or motor variance (Fig 2d, f). Moreover, in Exp 3, the reaction time priming effect was similar after both Opposite and Standard trials (Fig S4e), suggesting the effect may be related to target detection rather than reaching, per se.

### Encoding noise enhances the repulsive sequential bias

Another key prediction of the efficient coding theory is that the sequential bias should be modulated by encoding noise. Specifically, increased noise within the system should lead to a broader distribution of the response signal. Consequently, the average response of the system will exhibit a larger repulsive bias relative to the prior mean (Fig 4a). To examine this, we compared the sequential bias in reaching movements between the dominant hand and the non-dominant hand, with the premise that encoding a movement with the non-dominant hand incorporates more noise (Fig 4b), which should increase the repulsive bias. In Exp 4, we performed a similar center-out reaching task and participants were randomly instructed to use one hand per trial. To make sure that participants adhered to the instruction, we performed this experiment in a lab setup with the experimenter supervising the task (Fig 4c). Participants moved their occluded hand on a tablet, with the visual stimulus displayed on a monitor directly above (Fig 4c). Consistent with our assumption, participants showed significantly lower motor variability with the dominant hand compared to the non-dominant hand (*t*(23)=4.9, *p*<0.001, d=1.0; Fig 4d).

**Figure 4.**
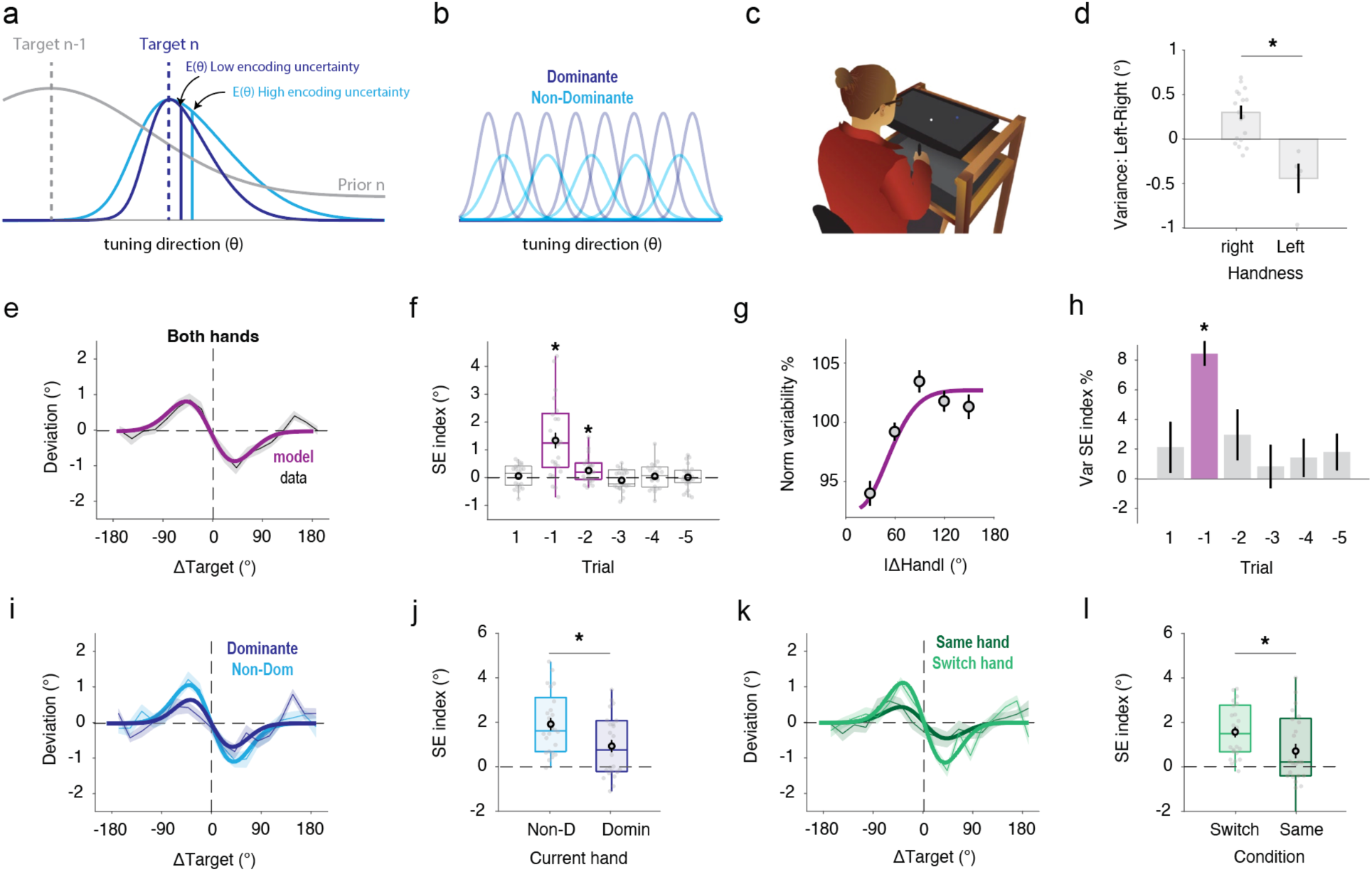
The repulsive sequential bias increased with the encoding noise in the motor system. a) Illustration of how encoding noise increases the repulsive bias based on the efficient coding model. b) We assumed that the non-dominant hand has fewer movement planning units with higher encoding uncertainty, indicated by a broader tuning profile. c) Illustration of the lab-based setup. d) Difference in motor variance between the left and right hand. Right-handed participants showed nosier movements using their left hand and vice versa. e-f) The sequential effect of motor bias (e) and the SE index (f) when data from both hands were collapsed. g-h) Sequential effect of motor variance (g) and (h) the estimated coefficients from a general linear model. i-j) A stronger repulsive bias was observed when the current movement was performed using the non-dominant hand compared to the dominant hand. k-l) A stronger repulsive bias was observed when participants switched hands compared to the situation when they used the same hand for trial N-1 and trial N. Error bars and shaded areas indicate standard error. *, p<.001.

The results from Exp 4 provide compelling support for the efficient coding model. All main results were consistent with what we had observed in the online experiments. When we combined the data across the two hands, we observed a repulsive sequential bias (N-1: *t*(23)=4.7, *p*<.001, d=0.96; N-2: *t*(23)=2.6, p=.02, d=0.55; Fig 4e-f) and increased movement variance (N-1: *p*<.001, Bootstrap, Fig 4g-h). Importantly, we observed a larger sequential bias when the current movement was performed using the non-dominant hand compared to the dominant hand (t(23)=4.7, p<0.001, d=0.97; Fig 4i-j), supporting the prediction of the efficient coding theory that bias escalates with encoding noise in the motor system. Interestingly, we also observed a larger sequential bias when participants switched their hands (namely using different hands in trial N and N-1) compared to using the same hand (t(23)=6.7, p<0.001, d=1.4; Fig 4k-l). This effect is likely due to dynamic allocation of encoding resources across hands, leading to increased encoding variability immediately after a hand switch and, consequently, a larger bias. Alternatively, more units might be recruited when repeating a movement with one hand, reducing encoding noise as well as the sequential bias.

To further examine how the variability in the motor system influences the sequential bias, we analyzed the correlation between motor variance and the sequential effect (SE) index in Exp 4. Consistent with the prediction of the efficient coding theory, we found a positive correlation between individual differences in variance and the SE index for both hands (Fig 5a). A positive correlation was also found when we re-examined the data from Exp 1 and 3 (Fig 5b-c). Those results together further supported the hypothesis that motor planning follows the principles of efficient coding.

**Figure 5.**
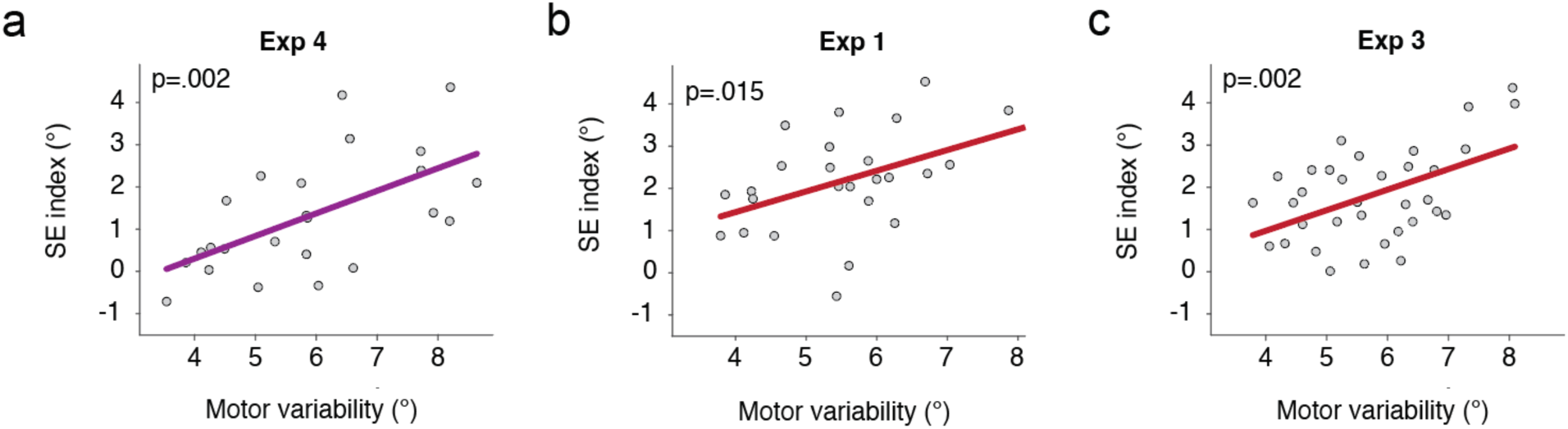
The repulsive sequential bias increased with the motor variability across participants. Correlation between the SE index and the motor variance for Exp 4 (a), Exp 1 (b), and Exp 3 (c). Each dot is one participant. The colored line shows the best-fitted linear model. The p-value was measured from the Pearson correlation.

## Discussion

The nervous system is dynamically tuned based on past experiences to be optimally prepared for the current task. These adaptive processes induce sequential effects in behavior^38^. In the present study, we examined the computational principles that govern the sequential effects in motor planning using reaching tasks with targets randomly sampled from a uniform circular distribution. Contrary to the prevailing belief that motor control is predominantly governed by Bayesian principles^3,9,10,45^, we demonstrated a novel repulsive sequential bias in which the current movement is biased away from the direction of the previous movement. This anti-Bayesian behavior suggests that, in less predictable environments, an alternative mechanism may also exist in motor control.

Interestingly, the sequential effects we observed indicate that motor planning follows the efficient coding model, a theory that posits the nervous system maximizes mutual information by dynamically allocating resources to follow the prior distribution^25,27,28^. Efficient coding has been widely applied to perceptual systems^27,33,34,46^, and our results extend these principles to motor control. Our results confirm three key predictions of the efficient coding model in motor planning. First, the sequential bias was repulsive. Second, this effect was amplified when reaching variance increased, as evidenced by differences between the left and right hands and the strong effect of individual differences in motor variability. Third, motor variance also exhibited a sequential effect, with movements closer to the previous one showing lower variability and higher accuracy. Together, these sequential effects in reaching would not result from simple fatigue. Instead, they reflect an adaptive mechanism that enhances motor accuracy.

The temporal dynamics of the sequential effect highlight the flexibility of the motor system in adapting to environmental statistics. Remarkably, the repulsive effect diminished after only three trials (or ∼30s based on Exp S1), suggesting that priors for efficient coding might be maintained by motor working memory^47^. This adaptability is likely to be ideal for motor planning, since one may need to repeat some quick actions, such as pressing a buzzer or knocking on a door. In contrast, it is less common for identical reaching movements to be necessary after a lapse of a minute. Consistent with this notion, in Exp 4, we found that the sequential effect decreased when participants consistently used one hand, as opposed to alternating between hands. This observation suggests that encoding resources can be preferentially allocated to the active hand when repeating the movement with the same hand, thereby increasing motor accuracy.

Sequential effects are well-established in the visual system^37,48,49^, but our results clearly distinguish current motor effects from the visual serial dependence reported in previous studies. First, the direction of the motor sequential bias observed in our study is opposite to the serial dependence typically seen in visual tasks^37,41,43,49^. In our motor tasks, the current movement is repelled from the previous movement, whereas in visual perception tasks, the perceived target is usually attracted toward the previous target position^41^. Second, when perception and movement are in opposite directions (Exp 4), sequential effects are determined by the movement direction rather than the target position^43^. Third, our sequential effects are significantly modulated by handedness, further suggesting that they are associated with movement rather than perception.

The opposite directions of sequential biases in the motor and visual systems may reflect their distinct functional purposes and underlying mechanisms. In visual perception, the physical world is continuous and relatively stable^50–52^, making it crucial for the visual system to maintain a consistent representation of the environment^39^. Thus, employing Bayesian decoding—which induces an attractive effect—is beneficial for visual perception, ensuring a coherent and reliable perception of the environment^40,53^.

While the objects of perception in the world are physically stable, the actions we need to make upon these objects can change from moment to moment. The motor system, therefore, operates under distinct constraints. As motor goals can vary quickly over time^20^, an attractive bias may not benefit motor planning in such dynamic contexts, whereas a decorrelation mechanism, such as efficient coding, may help to minimize repetitive errors and counteract attractive biases from other sources. Additionally, Bayesian integration is typically associated with decoding processes that interpret sensory inputs based on prior knowledge^27,29^, whereas motor planning, at least by definition, is an encoding rather than decoding process, making Bayesian decoding mechanisms less applicable.

An open question remains whether the repulsive motor bias and the attractive biases induced by visual serial dependence may counterbalance each other. To isolate motor effects, we designed our study to minimize potential visual influences. For instance, visual effects are usually linked to perceptual uncertainty^27,54,55^. A typical visual task for serial dependence usually features vague targets, brief presentations, or disappearance upon response. In contrast, our study used high-contrast targets that remained visible during movement. Future studies could employ motor tasks with higher perceptual uncertainty, allowing both visual serial dependence and motor sequential effects to be observed. It is possible that the strengths of visual and motor effects are correlated and thus counteract each other.

Interestingly, the sequential bias identified in our study contrasts with a history-influenced effect known as use-dependent learning in motor control. Use-dependent learning refers to a bias toward repeated movements in the same direction^15,16,56^, aligned with predictions of a Bayesian model. However, use-dependent learning only occurs for relatively predictable targets, whereas targets in our experiment were completely random. Moreover, the time course of use-dependent learning is very different from the repulsive sequential bias we observed. Use-dependent learning typically requires multiple trials to manifest^15,19,22^, whereas the current repulsive effect emerges after a single trial. Finally, use dependent learning usually last for tens of trials while the repulsive sequential effect reported here lasted for only two trials.

Our results suggest that the differing timescales of use-dependent learning and efficient coding may reflect a progressive shift of motor control strategy in developing a skilled movement. The repulsive bias induced by efficient coding could be beneficial in a volatile environment for several reasons, such as reducing repeated errors and enhancing sensitivity to changes in the environment, body, or target^57,58^. However, there are instances where repeated, stable movement is desirable, such as when hammering a nail while holding it steady with one’s fingers. Initially, efficient coding may increase the precision of movements. Those movements might be slow and performed with caution so that the system can correct for undesired exploratory errors with online control. As the activity continues, use-dependent learning dominates to stabilize the movement, facilitating smooth and stable repetition without conscious control. Future studies should explore the intriguing possibility that there is a transition between efficient coding and use-dependent learning, which may reveal how the system balances the competing needs of flexibility and stability in motor planning.

## Methods

### Participants

Testing was conducted online for Exp 1, 2, 3, S1 and in the lab for Exp 4. For the online studies, 154 young adults (76 female, age: 26.7 ± 4.9 y) were recruited using the Prolific.io. The participants performed the experiment on their personal computers through a web-based platform for motor learning experiments. Based on a prescreening survey employed by Prolific, the participants were right-handed and had normal or corrected-to-normal visions. These participants were paid $8/h. For the lab-based experiments, we recruited 24 undergraduate students (15 female, mean age = 21.42y, SD = 3.78y) from the University of California, Berkeley community. 19 of the participants were right-handed and 5 of them were left-handed based on their scores on the Edinburgh handedness test^59^ and had normal or corrected-to-normal vision. These participants were paid $20/h. All experimental protocols were approved by the Institutional Review Board at the University of California, Berkeley. Informed consent was obtained from all participants.

### Design and procedure

#### Experiment 1

Exp 1-3 were performed using our web-based experimental platform^60,61^. The code was written in JavaScript and presented via Google Chrome, designed to run on any laptop computer. Visual stimuli were presented on the laptop monitor and movements were produced on the trackpad. Data was collected and stored using Google Firebase.

26 participants (20 females) took part in Exp 1. To start each trial, the participant moved the cursor to a white circle (radius: 1% of the screen height) positioned in the center of the screen. After 500 ms, a blue target circle (radius: 1% of the screen height) appeared with the radial distance set to 40% of the screen size. Target locations were randomly generated from 1°-360° with a minimum step of 1°. The participant was instructed to produce a rapid shooting movement, attempting to move through the target. A feedback cursor (radius: 0.6% of screen height) appeared for 100 ms when the amplitude of the movement reached the target distance, indicating the angular position of the hand at that distance. The feedback cursor and target were then extinguished. If the movement time was >300 ms, the message “Too Slow” was presented on the screen for 500ms. At the end of the trial, the position of the cursor was reset to a random position within a circle centered at the start position with a radius 4% of the target distance. The participant moved the cursor back to the start position to initiate the next trial. Each participant completed 1080 trials in total.

#### Experiment S1

We aimed to examine the temporal modulation of serial dependence in Exp 3. Compared to Exp 1, we extended the inter-trial interval to either 6s (n = 28, 10 females) or 18s (n = 23, 12 females) for two groups of participants, respectively. A message “wait” would be presented on the monitor between two trails. Participants were instructed to put their right hand on the trackpad and rest until they saw the message “move to center” which indicated the start of a new trial. Participant completed 880 (6s condition) or 360 trials (18s condition).

#### Experiment 2

To confirm the effect observed in Exp 1a was not due to the existence of endpoint feedback, we replicated Exp 1 in 2 without presenting any feedback after the movement. Other details of Exp 2 were identical to Exp 1. 36 participants (21 females) took part in Exp 2.

#### Experiment 3

Exp 3 was designed to examine whether the sequential effect was induced by perception or movement. 41 participants (13 females) took part in Exp 4. The procedure of Exp 3 was essentially the same as in Exp 1. To evaluate whether this repulsive effect was perception-based or motor-based, we included trials (25%) in which the participants were instructed to move in the opposite direction of the target. Before each trial, an instruction message would appear on the screen to instruct participants to either “move to target” or “move to opposite.” There were no consecutive “opposite” trails. Each participant completed 960 trials in total.

#### Experiment 4

Exp 4 was designed to examine how motor variability influences the sequential reaching bias. 24 participants (15 females, 19 right-handed and 5 left-handed) performed the experiment in the lab setup. Participants performed a center-out reaching task, holding a digitizing pen in the right or left hand to make horizontal movements on a digitizing tablet (49.3cm x 32.7cm, sampling rate= 100 Hz; Wacom, Vancouver, WA). The stimuli were displayed on a 120 Hz, 17-in. monitor (Planar Systems, Hillsboro, OR), which was mounted horizontally above the tablet (25 cm), to preclude vision of the limb. The experiment was controlled by custom software coded in MATLAB (The MathWorks, Natick, MA), using Psychtoolbox extensions, and ran on a Dell OptiPlex 7040 computer (Dell, Round Rock, TX) with Windows 7 operating system (Microsoft Co., Redmond, WA).

Participants made reaches from the center of the workspace to targets positioned at a radial distance of 8 cm. The start position and target location were indicated by a white annulus (1.2 cm diameter) and a filled blue circle (1.6 cm), respectively. Vision of the hand was occluded by the monitor, and the lights were extinguished in the room to minimize peripheral vision of the arm. Feedback, when provided, was in the form of a 4 mm white cursor that appeared on the computer monitor, aligned with the position of the digitizing pen.

To start each trial, a letter “R” or “L” would be presented within the start circle to inform participant which hand to use on this trial. Participants used the instructed hand to hold the pen and put the other hand on the side. The experimenter supervised the whole experiment to make sure the participant applied the correct hand. After maintaining the cursor within the start circle for 500 ms, a target appeared. The participant was instructed to make a rapid slicing movement through the target. Right after their movement amplitude reached 8 cm, a cursor would be presented at that position for 1 s, providing feedback of the accuracy of the movement (angular position with respect to the target). After this interval, the target and cursor were extinguished. Another letter appeared at the start position and participants changed hand accordingly. To guide the participant back to the start position without providing angular information about hand position, a white ring appeared denoting the participant’s radial distance from the start position. Once the participant moved within 2 cm of the start position, the ring was extinguished, and a veridical cursor appeared to allow the participant to move their hand to the start position. If the movement time was >300 ms, the audio “Too Slow” was played right after the reaching.

The required hand was pseudorandomized so there were 4 left-hand and 4 right-hand trials within every 8 trials. Target locations were randomized in a way that both hands would visit targets from 1°-360° (with a step of 1°) within every 720 trials. The whole experiment included 2160 trials and took about 4 h. However, we allowed participants to end the experiment based on their convenience. 14 out of 24 participants finished all trials; other participants finished 1000∼2000 trials.

### Data Analysis

We calculated the error of hand angle as a difference between the hand position when it reached the target distance and the target position. A positive hand angle denoted that the hand position was more clockwise than the target at the target radius. Trials with a movement duration longer than 500 ms or an error larger than 60° were excluded from the analyses. For the web-based experiment, 1.5% trials were removed. For the lab-based experiment, 0.4% of trials were removed.

To analyze the sequential effect in movement, we regressed out the influence of systematic bias. Specifically, we fitted a function between motor bias and target angle using a polynomial function with a maximal power of 10 for each participant. We then subtracted this motor bias function from the motor error. The residual error was defined as motor “deviation” and applied to analyze the sequential effect^35,36,41,42^.

To analyze the sequential bias of the movement direction, we measured the function of how motor deviation changed as a function of the difference of target position between trial N-1 and trial N (defined as ΔTarget). A function lies in Quadrants II and IV will suggest that the movement error was in the opposite direction of the previous target. To quantify sequential bias, we introduced the SE index that takes the difference between the average error within −90°-0° Δ Target and the average error within 0°-90° Δ Target. A positive SE index means a repulsive sequential effect and vice versa.

To further examine how the variability in the motor system influences the sequential bias, we calculated the Pearson correlation between motor variance and the SE index, for Exp 1, Exp 3, and Exp 4 (only for trails after a standard trial). The motor variance was very large in the no-feedback condition of Exp 2 compared to all other experiments, likely because of the drifting sensorimotor map without visual calibration^62,63^. As such, the motor variance in Exp 2 was not a good measurement of encoding variability.

To analyze the sequential effect in motor variance, we calculated the absolute difference between the movement N-1 and movement N (|Δ Hand|). We flipped the sign of the deviation if Δ Hand was negative. Then we calculated the variance of deviation within each bin of 30°. The variance was then normalized by the average the average variance for each participant. To examine the tendency of how variance changed as a function of |Δ Hand|, we applied a general linear model:

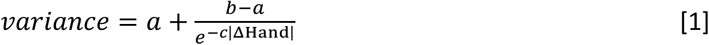

where a, b, c are the three free parameters. The SE index was defined as the change in the output of the function when |Δ Hand| increases from 30° to 150°. To estimate the distribution of the SE index, we applied bootstrap resampling 1000 times.

To analyze the priming effect in the reaction direction, we normalized the reaction time (RT) by subtracting the average reaction of each participant. We then plotted a function of how the normalized RT changes as a function of |Δ Target|. The RT SE index is the difference between the average normalized RT within 180°-90° |Δ Target| and the average RT within 0°-90° |Δ Target|. Simple t-tests were conducted to determine whether the SE indexes were significantly different from 0. We confirmed that the data met the assumptions of a Gaussian distribution and homoscedasticity for all tests. The significance level was set at p < 0.05 (two-tailed).

## Model

### Efficient coding model

We assumed that the motor system encodes a movement direction (*m*) based on an observed target direction (*θ*) following the rule of efficient coding. This model is based on previous models of efficient coding in perception^26,27,29^. A key assumption of the model is that the encoding system allocates its resources to maximize the mutual information *I*[*θ*,*m*] between input *θ* and output *m*. By imposing a constraint to bound the total coding resources of the system^27^, this requires the Fisher information *J*(*θ*) to be matched to the stimulus prior distribution (*p*(*θ*)): *p*(*θ*)∝*J*(*θ*). As such, coding resources are allocated such that the most likely movement direction is coded with the highest accuracy.

We next calculated the likelihood functions of how the system responds to different target directions with constraints of the prior distribution. Technically, the likelihood functions can be computed by assuming a symmetric Gaussian noise structure in a space where the Fisher information is uniform (the motor space, 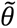), and then transforming those symmetric likelihood functions back to the target space (*θ*). To construct a motor space with uniform Fisher information based on the prior distribution of *θ*, one defines a mapping *F* from the target space (*θ*) to the motor space 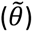, following^27^:

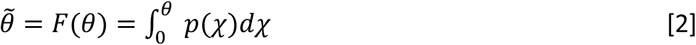

Given an input *θ*, the output *m* is computed as follows^26^. We first calculated the response value *r*, which would be of the form *θ* + *ϵ *, where *ϵ * represents an error due to the intrinsic encoding noise of the system. Note that *ϵ * follows an asymmetric distribution in the target space (Fig 1b). Let 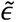 represent the transformation of *ϵ * to the motor space. Since we were assuming 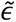 has a symmetric Gaussian distribution, the response value in the motor space would be 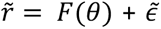, where 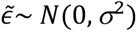; which gives a response value in the target space of:

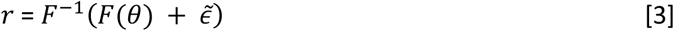

Next, we assumed that the system knows that its response *r* is noisy. Therefore, it generates a distribution of the form *r* + *δ*, where *δ* follows the same distribution as *ϵ *. If 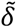 represents the transformation of *δ* to the motor space, this distribution will be of the form *F*^-1(^(*r* + *δ*), where 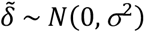. Finally, the system returns the expected value of this distribution as the output *m*:

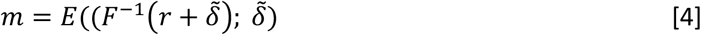

where *E*(X; z) means the expected X as z varies. We defined 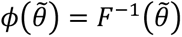. In the small-noise limit, we can take a second-order Taylor expansion:

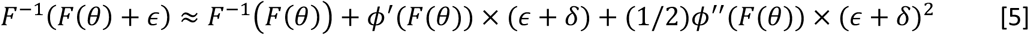

Considering [3]-[5] together, we have:

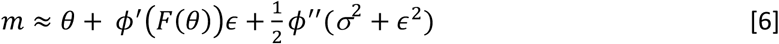

To estimate the motor bias predicted by the model, we calculated the expected value of 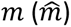 when *ϵ * varies can be expressed as:

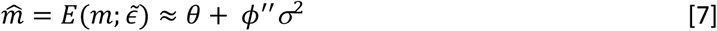

And the variance of *m* across trials can be expressed as:

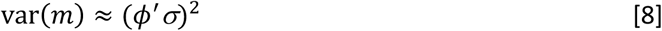

Since the sequential effect in movement is influenced principally by the last movement, we assumed the prior of the motor planning system is a mix of a uniform distribution across the whole space and a Gaussian distribution centered at the last target direction (θ*_n_*_-1_):

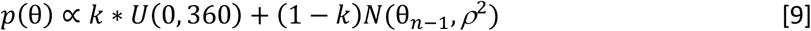

where *k* is a scale factor controlling the relative contribution of the two distributions; *ρ* indicates the width of influence from the previous trial. To simulate the bias and the variance predicted by this efficient coding model, we computed the numerical approximation of *ϕ*” and *ϕ*’ based on this prior function using an incremental approach.

This model includes three free parameters: *k* and *ρ*, which determine the shape of the prior, and *σ* which reflects the encoding noise. Since many parameter sets can produce similar bias patterns but different variance functions, we fitted the model by constraining both the bias and variance functions. Specifically, we predicted the deviations using the model and calculated the mean squared error (MSE) between these predictions and the observed data across all trials and participants. Additionally, we computed the variance function and calculated the MSE between the predicted function and the empirical group-level variance. We then minimized the sum of these two MSEs to determine the best-fitting parameters.

### Repeated Suppression model

We considered two alternative models to explain the sequential effects in the motor planning. The first model is a repeated suppression model, which assumes that neurons tuned to a specific direction become less sensitive after repeating a similar movement. Those modulations can enhance the sensitivity to the changes in the environment or/and encourage exploration. Here we applied a population coding model with a group of neurons with Gaussian-shaped tunning functions. For a target direction θ, the unit tunned to *i* (*i* ∈ [0, *π*]) direction generates a response *r_i_* as follow:

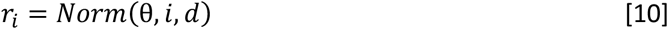

where *Norm*(θ, *i*, *d*) is the probability density function of a Gaussian distribution with a mean of *i* and standard deviation of *d*. The output of the system is determined by summing the activation of all neurons:

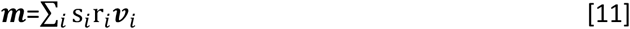

where ***v**_i_* is a vector representing the tuning direction of unit *i*, ***m*** is a vector pointing towards the movement direction, and s*_i_* is the sensitivity of unit *i*. After a movement in trial *n*, s*_i_* is updated based on the strength of the activation in unit *i*:

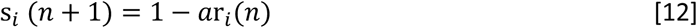

where *a* ∈ [0,1] is the suppression rate. As such, units that response more to the target in trial *n* will be more suppressed in the next trial.

### Bayesian Decoding model

The second alternative model we applied is a classic Bayesian Decoding model that utilizes the prior distribution of *θ* to improve performance^3,64,65^. The system generates a response *r* based on a target direction *θ*. Considering Gaussian encoding noise, the relationship between *r* and θ can be expressed as follows:

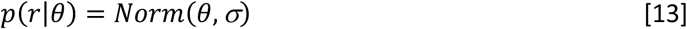

Importantly, the model assumes that the system utilizes both the prior and this likelihood function to form a posterior estimation following Bayesian rules:

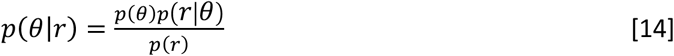

where *p*(*r*) is a constant; *p*(*θ*) is the prior; *p*(*r*|*θ*) is the likelihood; and *p*(*θ*|*r*) is the posterior. The output of the system (*m*) is the posterior mean. For the Bayesian model, we used the same prior distribution (see [9]) as the efficient coding model.

## Acknowledgments

We thank Richard Ivry for helpful discussions.

## Data availability

All data and code are available at https://github.com/shion707/MotorEC.

## Supplementary Information

### Supplementary Result 1: Temporal dynamics of the sequential effects in Reaching

In Exp 1-2, we observed a repulsive effect only from the 1-back and 2-back trials. This motivated us to further examine the temporal regulation of the repulsive sequential effect. We tested whether the rapid drop in the sequential effect solely depended on time, or whether it also depended on the number of trials. As such, in Exp S1, we extended the inter-trial interval (ITI) to either 6s or 18s for two groups of participants and compared the results with Exp 1 (0s ITI) (Fig S1). While the strength of the sequential effects decreased with time, both the repulsive bias (*t*(23)=5.2, *p*<0.001) and the variance modulation (*p*=0.020) could be observed for trial N-1 if participants simply wait for 6s between two trials. This result contrasted with the N-3 trial in Exp 1, which was ∼6s prior to the current movement (Fig S1b, d), but no sequential was observed. Moreover, the repulsive bias remained significant even in the 18s ITI condition (Fig S1b). The SE index of the motor variance in the 18s ITI conditions showed a positive trend but did not reach significance (*p*=0.16, Fig S1d). These results suggested that the attenuation of the sequential effects depended on both passing time and intervening information.

**Figure S1.**
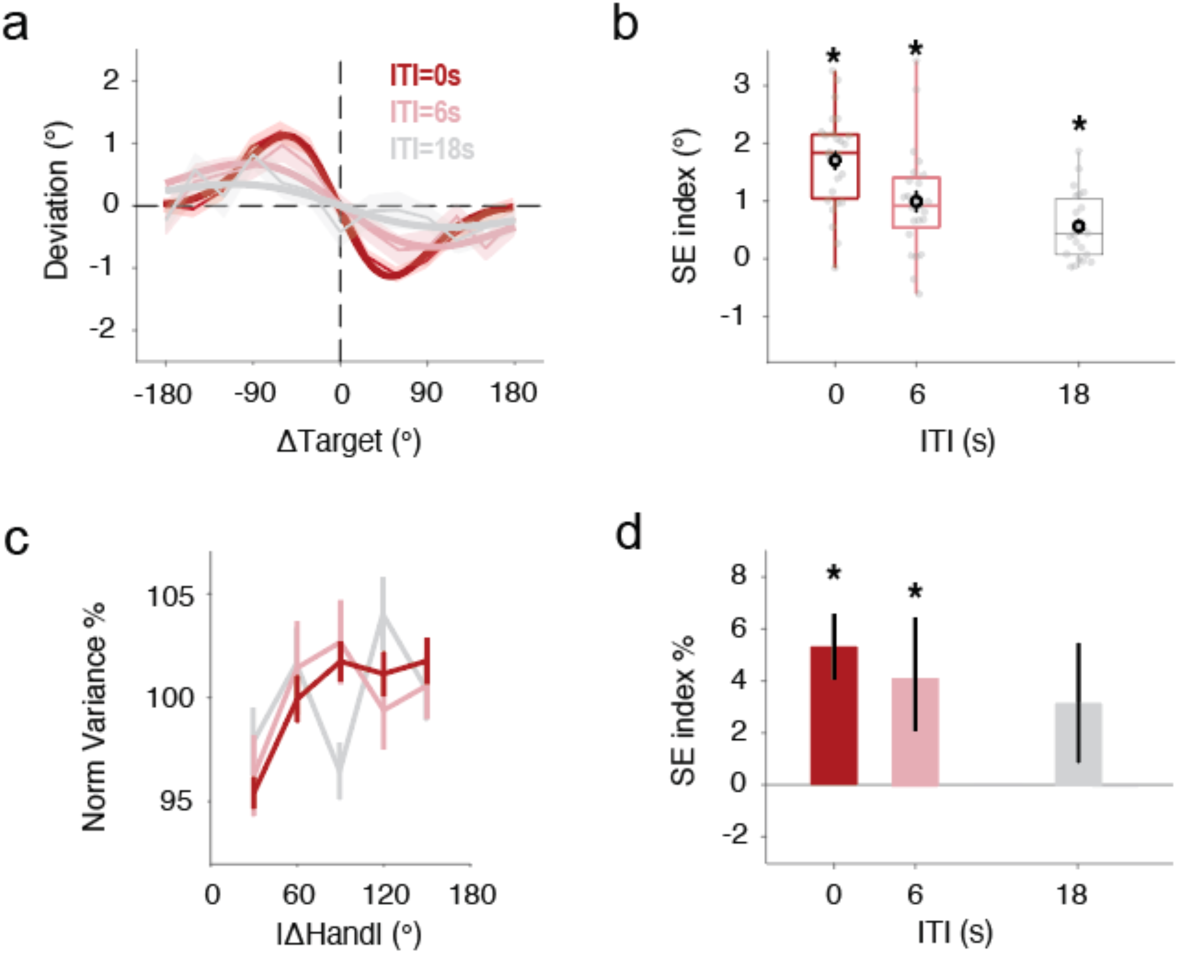
Temporal dynamics of the Sequential effects in reaching. (a) Sequential effect in movement direction for three ITI conditions. The thinner lines indicate the data, and the thicker lines indicate the prediction of the efficient coding model. (b) SE index with different ITI. Different from what has been observed in the 3-back condition in Exp 1 (which had a ∼6s delay), the SE index was significant in the 6s and 18s ITI conditions, indicating the effect decreased with both time and trial number. Error bar and shad area indicate standard error. (c-d) Sequential effect in motor variance was modulated by ITI. A significant SE index was found in the 6s ITI condition. *, p<.02.

**Figure S2.**
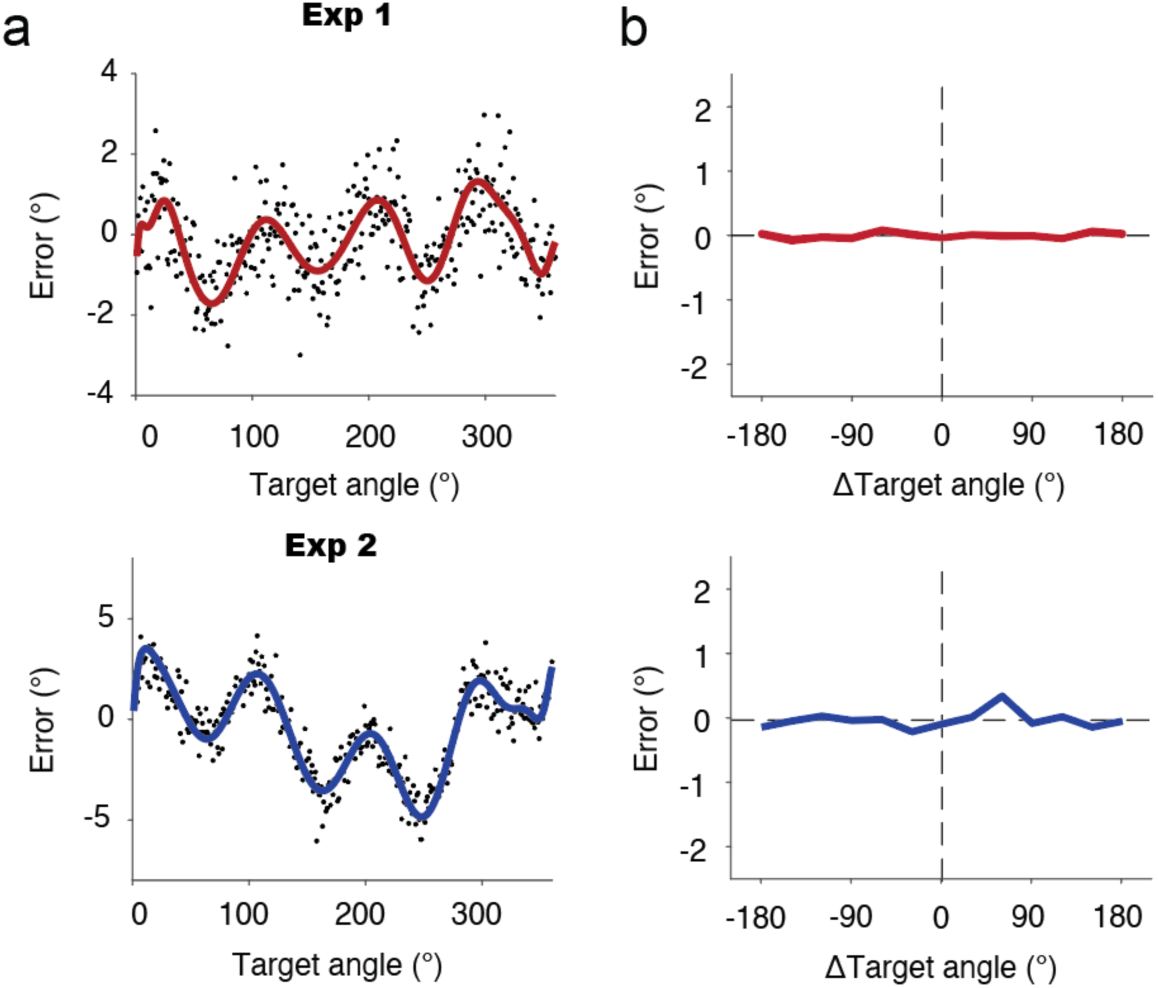
Systematic motor biases did not generate sequential effects. a) Systematic motor bias relative to target position in Exp 1 (upper panel) and 2 (lower panel). Participants exhibited greater bias in Exp 2, where feedback was absent. Black dots indicate data. The thick line represents a smoothed function, obtained by fitting a polynomial function with a maximum degree of 20. b) Simulation of sequential bias based on the motor bias function and the sequence of targets presented to participants. The systematic motor bias did not generate any sequential effect.

**Figure S3.**
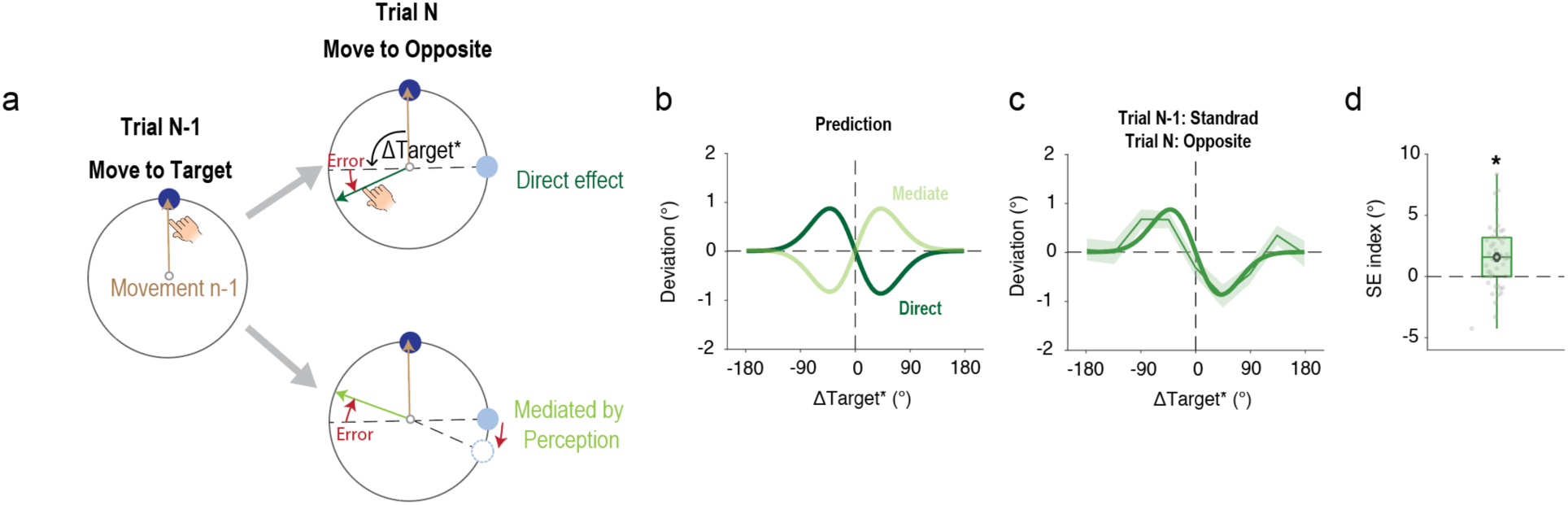
The current movement was directly influenced by the previous movement, with no mediation of perception. a) When trial N was an Opposite trial and trial N-1 was a Standard trial, it allowed for examination of whether the effect was directly caused by the previous movement repelling the current movement (direct hypothesis). Alternatively, it could be that the previous movement repelled the perception of the current target’s position, thus indirectly influencing the movement direction (mediated hypothesis). Here, Δ Target* was defined as the difference between the opposite position of the current target and the position of the previous target. b) Different predictions of the sequential bias following the direct hypothesis and mediated hypothesis, respectively. c) Sequential bias observed in Exp 3 was consistent with the direct hypothesis. The thin lines with shaded error bars indicate data, and the thick curve indicates the prediction of the efficient coding model. Shaded areas indicate standard error. d) SE index indicates that the current movement was significantly repelled away from the previous movement. *, p<.001.

**Figure S4.**
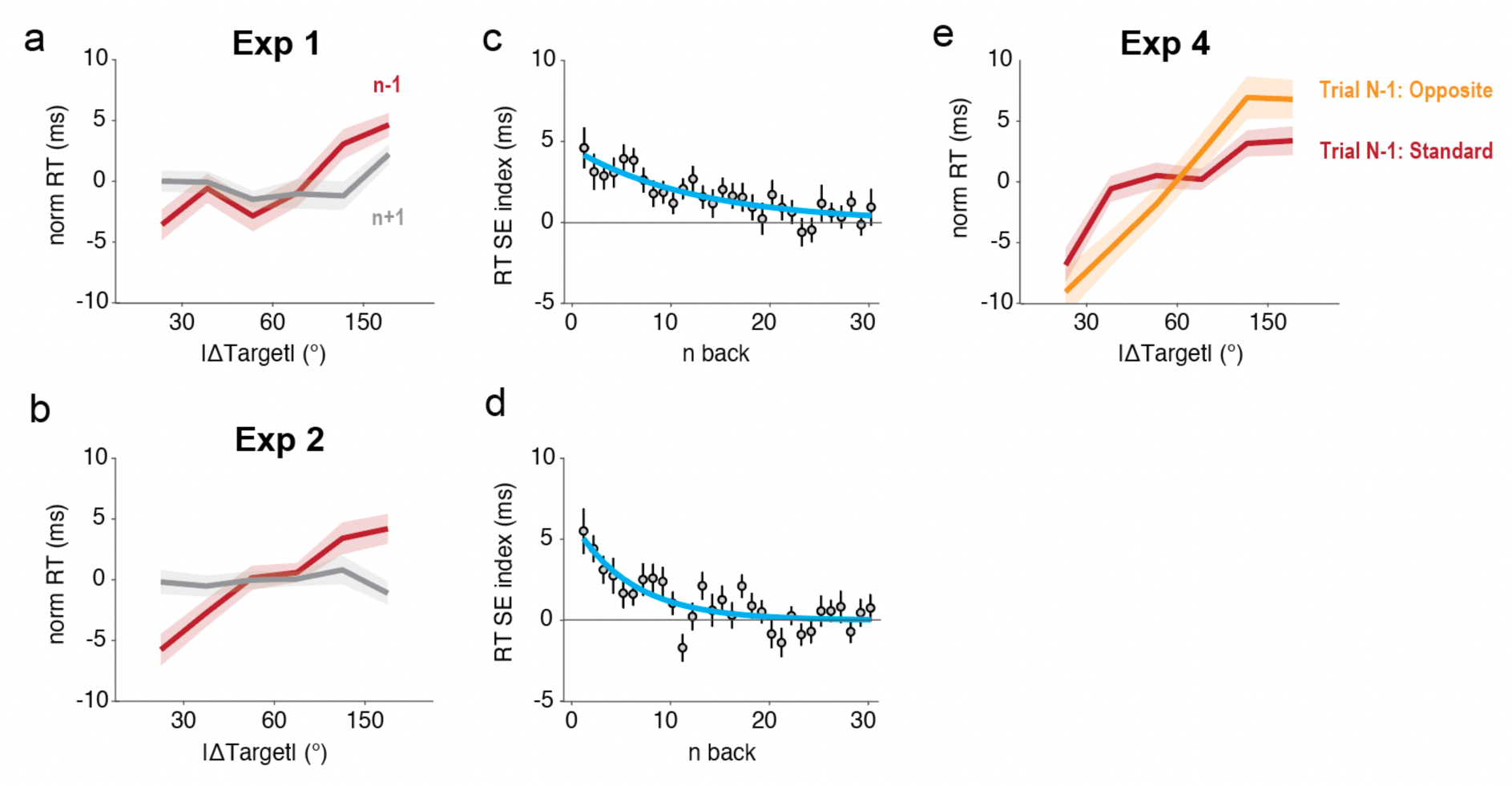
Priming of Reaction Times. a-b) Reaction time increased when the target in the previous (n-1) trial was further away from the current (n) trial in Exp 1-2. Reaction time was not correlated with the future trial (n+1). c-d) The SE index for reaction time lasted for more than 10 trials, differing from the temporal dynamics of sequential effects in movement direction or motor variance. e) The priming of reaction time was similar after an Opposite trial or a Standard trial in Exp 4, suggesting that the reaction time priming is mainly associated with previous target detection or localization.

